# Regional assessment of male murine bone exposes spatial heterogeneity in osteocyte lacunar volume associated with intracortical canals and regulation by VEGF

**DOI:** 10.1101/2023.02.08.527672

**Authors:** Jacob Trend, Alisha Sharma, Lysanne Michels, Patricia Goggin, Philipp Schneider, Katrin Deinhardt, Claire E Clarkin

**Author notes:** Corresponding author: Jacob Trend.

## Abstract

The porous bone cortex comprises an interconnected network of intracortical vascular canals and osteocyte lacunae, embedded within the bone mineral. Increases in cortical porosity reduce bone strength and increase fracture risk. To date, our understanding of mechanisms coupling the arrangements of the vascular: lacunar network in the bone cortex is poorly understood yet it could be key in establishing regulation of cortical porosity evident with age. Using synchrotron radiation-based computed tomography we develop automated tools to characterise the 3D spatial organisation and morphology of osteocyte lacunae, and the bone vasculature at the tibiofibular junction (TFJ), defining posterior, medial, lateral, and anterior regions in male C57BL/6 mice (n = 3). We also investigate the role of osteoblast-derived VEGF in regulating the 3D spatial arrangement by conditional disruption of VEGF in osteocalcin-expressing cells (OcnVEGFKO versus WT, n = 3). Regional lacunar phenotypes were assessed by 3D distance mapping of lacunar organisation surrounding the vascular compartments, including endosteal and periosteal surfaces, or intracortical canals. Surface-associated lacunae were indistinct in size across posterior, medial, lateral and anterior regions. However, lacunae associated with intracortical canals were significantly larger exclusively within the posterior region. In the absence of VEGF, the increased lacunar volume associated with posterior intracortical canals was lost. Our results suggest that the influence of intracortical canals on lacunar volumes is spatially regulated and sensitive to locally produced growth factors such as osteoblast-derived VEGF.

## Introduction

Degenerative bone diseases are characterised by bone fragility, with 76% of the reduction in bone strength in aged bone, caused by alterations in cortical porosity (McCalden et al. 1993). Bone porosity is organised into intracortical canals and osteocyte lacunae (Schneider et al. 2009; 2013), with intracortical canals housing the bone vasculature, providing the bone cortex with a blood supply, sustaining bone cell viability and growth (Bonewald 2011). Throughout life the bone vasculature retains specialised structural and functional roles, with endothelial cell heterogeneity existing at both the organ (Marcu et al. 2018) and tissue level (Ramasamy et al. 2014; Kusumbe, Ramasamy, and Adams 2014; Ramasamy et al. 2016; Xie et al. 2014), suggesting that specific lineages of the bone vasculature may provide specific functions. In bone, specific endothelial cell subtypes have been described and linked to angiogenic-osteogenic coupling (Ramasamy et al. 2014; 2014; 2016; Wang et al. 2017; Zhu et al. 2019). In addition to functional variability, the variable distribution of intracortical canals between bone regions has been documented in murine (Núñez et al. 2018; Schneider et al. 2007) and human (Perilli et al. 2015) bone. However, the function of region-specific arrangements of intracortical canals remains unclear.

The second component of cortical microstructure is the lacuna-canalicular network (LCN), an interconnected network of voids (lacunae) and canals (canaliculi) (Bonewald 2011; Goggin et al. 2016). Within the LCN resides the osteocyte network, which comprises osteocytes within lacunae and their interconnected cell processes. Osteocytes are terminally differentiated osteoblasts, which form the most abundant population of cells within bone (Bonewald 2011). Despite their abundance, our understanding of their regulation in health and disease remains poorly understood due to their isolation within the mineralised bone cortex. The situation of osteocytes within the mineralised bone cortex permits their primary functions as (1) mechanosensory cells (Tatsumi et al. 2007; Fritton and Weinbaum 2009) and (2) regulators of bone (re)modelling (Bonewald 2011), which may be initiated in response to stimuli including mechanical loading (Zhao et al. 2022) and lactation (Qing et al. 2012). Mechanical stress compresses the bone cortex, driving interstitial fluid flow through the LCN (Bonewald 2011). Mechanical strains within the LCN are transduced by osteocytes into biochemical cues driving bone resorption by osteoclasts and bone deposition by osteoblasts (Bakker et al. 2001).

In addition to initiating remodelling via osteoclasts and osteoblasts, osteocytes may remodel their local matrix through perilacunar remodelling (PLR), which allows osteocytes to remodel the bone matrix surrounding the LCN (Lane et al. 2006). During lactation, the local matrix is resorbed through PLR to sustain systemic mineral content, leading to a transient expansion of the LCN and enlargement of osteocyte lacunae (Qing et al. 2012). Importantly, Regional heterogeneity within the canalicular length, density and fluid flow is observed across murine bone (van Tol et al. 2020), suggesting that functional heterogeneity exists within the LCN in response to heterogeneous strain distribution described at this site (Núñez et al. 2018). However, similar to the description of heterogeneously distributed intracortical canals, an understanding of the heterogeneous morphology, distribution, and function of the LCN is not fully understood.

The proximity of osteocyte lacunae and intracortical canals within the bone matrix has prompted hypotheses that crosstalk exists between osteocytes and vasculature, acting to couple angiogenesis and osteogenesis. Intracortical canal networks may facilitate bone remodelling and development through interactions with osteoblasts and osteocytes (Goring et al. 2019) while their presence prevents the isolation of deeply embedded osteocytes (Núñez et al. 2018). However, while intracortical canals may influence osteocyte distribution, the mechanisms underpinning heterogeneity within the morphology and function of intracortical canals and osteocyte lacunae are undefined.

We have previously resolved the regional distribution of intracortical canals at the murine tibiofibular junction (TFJ) in the context of ageing (Núñez et al. 2018), in aged bone, intracortical canal density was reduced exclusively in the posterior region of the TFJ. As such, we have hypothesised that the study of the posterior TFJ may provide a unique, pathologically susceptible anatomical site to interrogate cortical bone microstructure. Importantly, when the bone cortex was not regionally analysed, the age-associated reduction in intracortical canal density was not observed, due to averaging across different cortical regions. While other studies have compartmentalised the bone cortex for morphometric analysis (Núñez et al. 2018; van Tol et al. 2020; Uniyal et al. 2021; Schneider et al. 2007; Cole et al. 2022; Chiba et al. 2013), the manual selection of regions may lead to inter- and intra-observer errors.

Although the assessment of the bone porosity remains challenging, due to its encasing within bone’s mineralised matrix, techniques such as synchrotron radiation-based computed tomography (SR CT) permit non-destructive, high-resolution imaging of bone in 3D (Schneider et al. 2007; Núñez et al. 2018; Goring et al. 2019). SR CT datasets may be segmented to isolate a negative imprint of the bone cortex (Schneider et al. 2004), permitting the study of cortical porosity and the networks within.

Driven by the hypothesis that porosity within bone is heterogeneous and governed by region-specific spatial cues, we here utilise SR CT to characterise intracortical canal and osteocyte lacunae distribution, morphology and spatial arrangements at the murine TFJ. In this work we describe (i) an automated means to segment SR CT scans of the bone cortex (ii), a 3D distance mapping technique to evaluate how osteocytes and the bone vasculature are spatially organised, and (iii) how a pathological phenotype leads to disruption of this vascular: lacunar organisation.

## Materials and methods

### Animals

Male C57BL/6 wild-type (WT) mice were bred in-house and maintained until 13 months of age. Post mortem, tibiae were dissected and fixed in 4% paraformaldehyde (pH 7.4 in PBS) for 48 h on rotation as described in (Núñez et al. 2018; Goring et al. 2019) and stored in 70% ethanol. Second-generation osteocalcin-specific VEGF knockout mice (OcnVEGFKO) were bred as described (Goring et al. 2019). Right tibiae were collected from 16-week-old male mice and prepared as above for 13-month-old C57BL/6 tibiae. The use of animal tissue was carried out in compliance with the Animals Act 1986.

### Synchrotron X-ray computed tomography

Synchrotron X-ray computed tomography (SR CT) was undertaken at Diamond Light Source (Harwell, UK) at Beamline I13-2 (Proposal 21843). In preparation for scanning, right tibae were dehydrated and mounted in paraffin wax to prevent sample movement during scanning. The TFJ was selected as the field of view and for each scan 2000 projections were acquired over a range of 360°. A photon energy of 18.5 keV was used with an exposure time of 500 ms, with a sample-detector distance of 15 mm. Images were acquired using the pco.4000 camera and imaged with an x4 objective lens to provide an isotropic voxel size of 1.65 µm and a cross-sectional FOV of 6.6 mm^2^. Datasets were corrected for ring artefacts and reconstructed using standard filtered back projection. The FOV was selected to incorporate a small region above the TFJ, with the first point of contact between the tibia and fibula defined as the beginning of the TFJ; 500 slices were selected from this site of first contact and used for analysis. The TFJ was set as the region of interest (ROI) to provide consistency in the analysed regions; further, the TFJ is an anatomical site consisting solely of cortical bone.

### Alignment of the tibiofibular junction

To standardise region separation across bone samples, the BoneJ moment of inertia function (Doube et al. 2010) was used to identify the longest axis of the TFJ (Fig 1B) and manipulated to match the image stack’s longest axis. This axis was rotated 45° and the dataset canvas size was adjusted to produce an equal height-to-width ratio – e.g., 4000 x 4000 pixels. Quadrants were identified via a selection of coordinates. For a canvas size of 4000*4000 pixels, quadrants of 2000*2000 were selected using the “SPECIFY” function. “MAKE INVERSE” was used to select the remaining 3 quadrants and “FILL” was used to fill these 3 quadrants with pixels with an intensity of 0 (black pixels, denoting empty space), leaving one quadrant with pixels of interest. Specifying the upper left quadrant (0-2000 in x, 0-2000 in y) isolated the posterior, the upper right quadrant (2000-4000 in x, 0-2000 in y) isolated the lateral, the lower left quadrant (0-2000 in x, 2000-4000 in y) the medial, and the lower right quadrant (2000-4000 in x, 2000-4000 in y) isolated the anterior region of the TFJ. This segmentation technique allocated 35.2 7% (± 1.49 %) to the posterior, 28.20 % (± 1.09 %) to the anterior region 16.67 % (± 0.23 %) to the lateral and 19.85 % (± 2.37 %) to the medial region.

**Figure 1.**
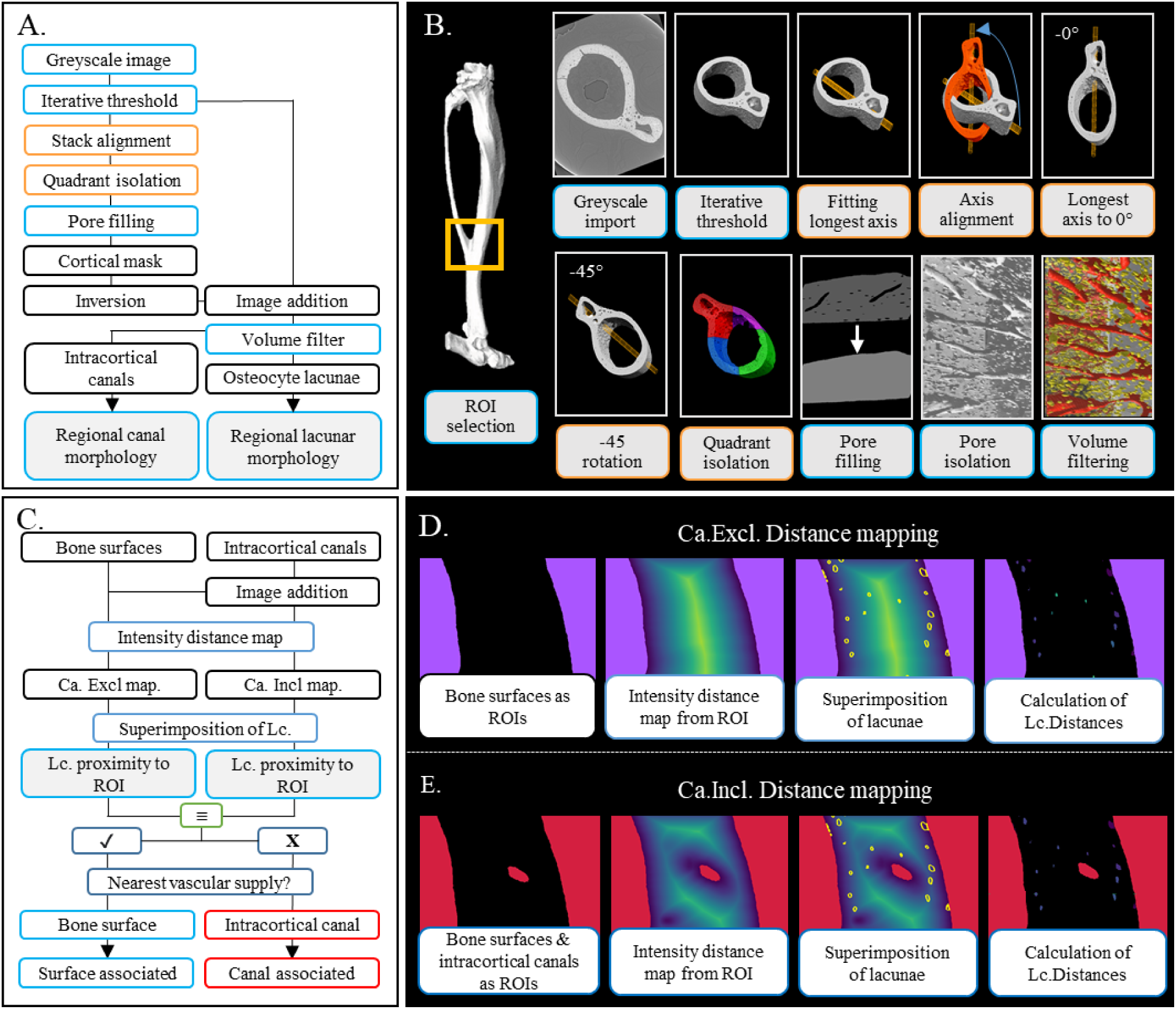
Image processing workflow for the automated regionalisation of the murine TFJ and the assessment of the microstructural organisation within. SR CT scans of the murine TFJ were imported into Fiji (A), and image stacks were aligned by fitting an ellipsoid to the TFJ, with the TFJ’s longest axis manipulated to 0°. The aligned TFJ was then split into quadrants. Regional cortical porosity was extracted and subjected to a volume filter to discard noise (0-25μm^3^) and to isolate osteocyte lacunae (25-2500 μm^3^) and intracortical canals (>2500 μm^3^). (B) Visualisation of image processing workflow described in (A). (C) Workflow for the assessment of osteocyte lacunae organisation surrounding bone surfaces and intracortical canals. Figures (D) and (E) visualise the workflow for Ca. Excl. distance mapping and Ca. Incl. distance mapping respectively.

### Extraction of cortical porosity

The binary-aligned TFJ image stacks were then duplicated and intracortical pores were filled with a series of binary dilate functions. Upon complete closure of intracortical pores, the cortical mask was eroded to return the mask to the size of the original cortex. The cortical mask was inverted and combined with the thresholded stack through image addition, yielding cortical porosity as described (Núñez et al. 2018; Schneider et al. 2007).

### Pore quantification

BoneJ’s particle analyser (Doube et al. 2010) was used to sort cortical porosity into noise (0 - 25 µm^3^), osteocyte lacunae (25 - 2500 µm^3^) and intracortical canals (> 2500 µm^3^). Cortical porosity (Ct. Po (%)), canal number density (N.Ca/Ct.TV (mm^-3^)), lacuna number density (N.Lc/Ct.TV (mm^-3^)), canal volume density (Ca.V/Ct.TV (%)), lacunar volume density (Lc.V/Ct.TV (%)), average canal volume (⟨Ca.V⟩ (μm^3^)), average lacunar volume (⟨Lc.V⟩ (μm^3^)), were calculated in concordance with standard nomenclature for bone morphometry (Núñez et al. 2018; Goring et al. 2019; Bouxsein et al. 2010; Schneider et al. 2007; Hemmatian et al. 2017). Lacunar volume distributions were visualised in Dragonfly. Osteocyte lacunae and intracortical canal containing image stacks were imported and converted into multi-ROI datasets so that each pore was treated as an individual region of interest. The object analysis tool was then used to quantify lacunar volume and mapped using the ‘warm metal’ LUT.

### 3D Lacunar distance mapping

Once osteocyte lacunae and intracortical canals were isolated, an image subtraction (binary cortical mask subtract binary intracortical canal image stacks) was used to create a cortical mask perforated by intracortical canals. This image was then inverted to convert the potential vascular surfaces (bone surfaces and intracortical canals) as the regions of interest and imported into Dragonfly. An intensity-based 32-bit distance map was then created from the inverted intracortical canal mask to map the distance of each pixel within the bone cortex, to the nearest surface of the inverted intracortical canal mask. Each pixel was then attributed a pixel intensity that directly relates to the distance from a vascular surface. Each increase in pixel intensity was indicative of a step size equal to the scan resolution (in this case, 1.65 µm, visualised in Fig 1C).

Next, lacunae image stacks were imported into Dragonfly and converted to a multi-ROI so that each lacuna was treated as a separate entity. Each lacuna was then superimposed onto the distance map, yielding a set of pixel intensities reflective of the pixels within that lacunae’s distance from a bone surface. The smallest of these distances was then isolated to yield ‘minimum lacunar distance’ and coloured using the ‘blue-green-yellow’ LUT. This was completed for each lacuna and averaged to yield the mean minimum lacunar distance (Mm.Lc.D).

### Lacunar association with vascular supply

To determine the role of intracortical canals in the spatial organisation of osteocyte lacunae and influence on Mm.Lc.D, the inverted cortical mask was also imported into Dragonfly and a distance map calculated between exclusively the endosteal and periosteal bone surfaces, in the absence of intracortical canals (Canal excluded distance mapping, termed ‘Ca. Excl’, shown in Fig 1D).

This was then compared to the lacunar distance mapping analysis including intracortical canals (Canal included distance mapping, termed ‘Ca. Incl’, Fig 1E). Should the minimum distance between each osteocyte lacunae and potential vascular supplies remain the same in both Ca. Incl and Ca. Excl analysis, then a bone surface is the nearest vascular supply. In this case, this lacuna is termed surface-associated. Should canal inclusion reduce the minimum lacunar distance, then intracortical canals reduce the distance between this individual osteocyte lacuna and potential vascular supplies, leading to classification as canal-associated. This analysis workflow may be visualised in Fig 1C.

This ‘association filter’ was applied to each lacuna in each region, sorting lacunae into canal-associated and surface-associated. From this, the percentage of lacunae associated with intracortical canals was calculated within each region and the distance was reduced by intracortical canals. In addition to Min.Lc.D, ⟨Lc.V⟩ was calculated for each lacuna.

### Statistical analysis

To investigate the role of region on bone’s microstructural characteristics in 13-month-old male mice, a one-way analysis of variance (ANOVA) was used. Tukey’s multiple comparisons were subsequently conducted to assess statistical differences between groups, as detailed in figure legends. For the assessment of cortical microstructural parameters in OcnVEGFKO, a paired two-way ANOVA was employed to compare littermates with subsequent Turkey’s comparisons used to compare the effect of OcnVEGFKO within each biological sex. Values are shown as mean ± SD, and p < 0.05 was deemed statistically significant. All statistical tests were performed in GraphPad Prism 6.0 (“GraphPad Prism Version 6.0”).

## Results

### Regional distribution of cortical porosity in the murine cortex

Microstructural analysis of the TFJ (Fig 2A) of 13-month-old male mice showed that global cortical porosity (Ct.Po%) was 1.38% (± 0.01%), in line with previous observations (Willey et al. 2008) of the TFJ in WT mice of a similar age. When assessed regionally, the posterior region possessed a significantly higher Ct. Po versus the anterior (+1.34%, p = 0.005), lateral (+1.08%, p = 0.017), and medial (0.93%, p = 0.036) regions (Fig. 2D, Table 1). Intracortical canal density was higher in the posterior region versus the anterior (+65.6%, p = 0.042), lateral (+71.7%, p = 0.030), and medial (+68.7%, p = 0.037) regions (Fig, 2E, Table 1). However, no significant regional differences were observed in canal volume density and average canal volume (Table 1).

**Figure 2.**
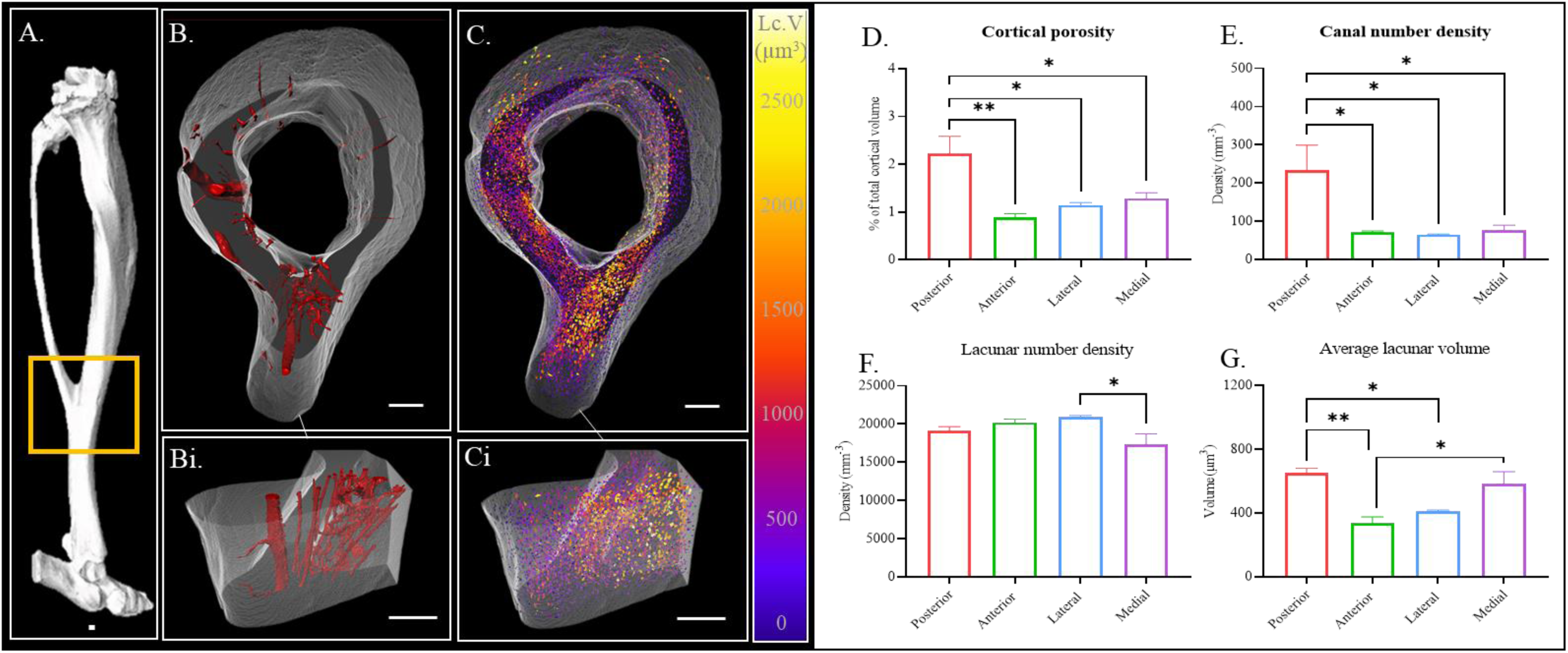
Regionalisation of murine cortical bone reveals region-specific microstructural properties. The regional microstructure was quantified at the TFJ (A) and separated into intracortical canals (B) and osteocyte lacunae, with colour mapping displaying osteocyte volume (C). The posterior TFJ is more porous than other regions (D), with intracortical canal number density (E) highest in the posterior TFJ (visualised in Bi). Isolation of osteocyte lacunae finds that although the differences in lacunar number density between regions are minimal (F), average lacunar volume varies between regions with the posterior TFJ possessing large lacunae as shown in (Ci) and quantified in (G). Large lacunae are coloured in yellow and orange. *Data shown as mean ± SD, * = p<0.05, ** p<0.01. Scale bars represent 200μm*.

**Table 1.** Regional microstructural morphometric measures for cortical regions within the murine TFJ.

| Parameter | Region |  |  |  | Anova |
| --- | --- | --- | --- | --- | --- |
| | Posterior | Anterior | Lateral | Medial | $p$-value |
| Ct.Po (%) | 2.22 ± 0.62 | 0.88 ± 0.13 | 1.14 ± 0.09 | 1.29 ± 0.19 | **0.005** |
| Ca.V/Ct.TV (%) | 0.97 ± 0.53 | 0.29 ± 0.064 | 0.29 ± 0.096 | 0.11 ± 0.03 | **0.050** |
| N.Ca/Ct.TV (mm) | 233 ± 92 | 76 ± 18 | 65 ± 2 | 71 ± 5 | **0.020** |
| $\langle \text{Ca.V} \rangle$ ($\mu\text{m}^3$) | 37765 ± 12526 | 40322 ± 10320 | 43674 ± 14626 | 16417 ± 5565 | 0.144 |
| Lc.V/Ct.TV (%) | 1.25 ± 0.13 | 0.59 ± 0.16 | 0.85 ± 0.020 | 1.18 ± 0.22 | 0.003 |
| N.Lc/Ct.TV (mm<sup>-3</sup>) | 19172 ± 685 | 17345 ± 1950 | 20901 ± 337 | 20235 ± 596 | **0.050** |
| $\langle \text{Lc.V} \rangle$ ($\mu\text{m}^3$) | 653 ± 47 | 339 ± 64 | 410 ± 15 | 585 ± 130 | **0.003** |

Osteocyte lacunar number density (N.Lc/Ct.TV, mm^3^, Fig. 2F, Table 1), average volume (⟨Lc.V⟩, Fig. 2G, Table 1) and volume density (Lc.V/Ct.TV, Table 1) were also calculated within each region of the TFJ. Lacunar number density was higher in the lateral region than in the anterior region (+ 19%, p = 0.046). Lacunae were larger in the posterior region than in the anterior (+ 48.1%, p = 0.004) and lateral (+ 37.2%, p = 0.019) regions, while medial lacunae were also larger than anterior lacunae (+ 42.1%, p = 0.180). Lc.V/Ct.TV was also higher in the posterior and medial regions than in the anterior region (+ 52.8% p = 0.003 and +50%, p = 0.007, respectively). The correlation of increased intracortical number density and large osteocyte lacunae within the posterior region motivated the use of 3D distance mapping techniques to assess the organisation of osteocyte lacunae surrounding intracortical canals between different regions.

### 3D distance mapping of osteocyte lacunae around intracortical canals

3D distance mapping was used to assess the organisation of osteocyte lacunae surrounding potential vascular supplies within bone; that is, endosteal or periosteal bone surfaces, or intracortical canals, within each cortical region. Global analysis revealed that the mean minimum lacunar distance (Mm.Lc.D) is (36.2 µm ± 0.99 µm), while regional analysis found no significant difference between regions (anterior 36.9 µm ± 4.7 µm, posterior 34.2 µm ± 1.8 µm, lateral 33.5 µm ± 2.3 µm, medial 40.4 µm ± 1.5 µm) as assessed by a one-way ANOVA (p = 0.139, Fig. 3C, grey). 49.3% (± 1.57%) of lacunae were localised within 25 µm of a bone surface or intracortical canal, while a minority of lacunae (6.4% ± 0.5%) were at a distance of >100 µm from a possible vascular surface.

**Figure 3.**
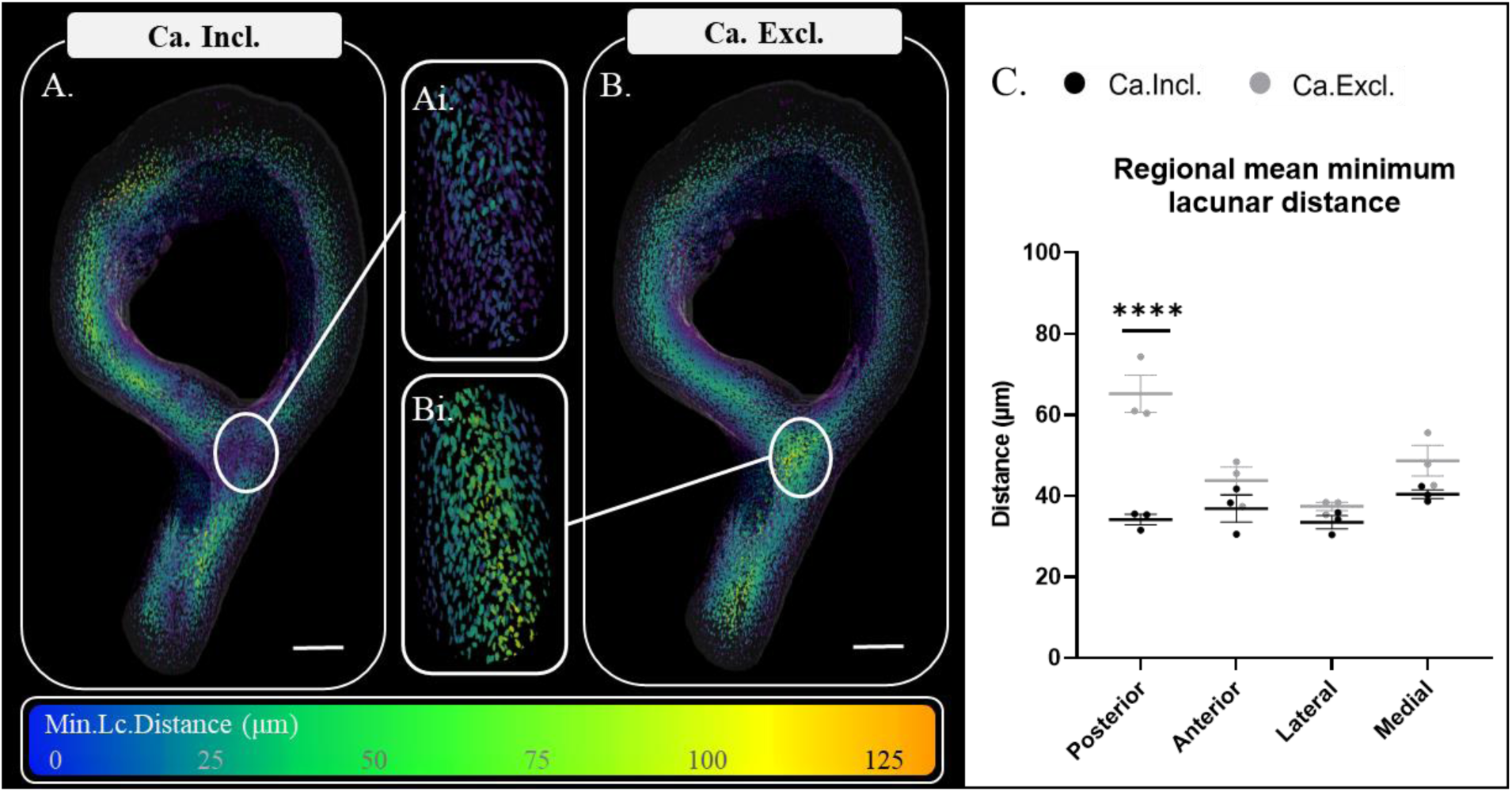
Posterior intracortical canals reduce the proportion of lacunae isolated deep within the cortex. Spatial mapping of mean minimum lacunar distance (Mm.Lc.D) from both bone surfaces and intracortical canals (B, Ca. Incl) and with canals excluded from the analysis (B, Ca. Excl). Removal of intracortical canals increased the Mm.Lc.D in the posterior region but not in any other region (C), with Ca. Incl maps shown in black and Ca. Excl maps in grey. Scale bars denote 200 μm. *Values reported are mean ± SD.* **** denotes a p-value < 0.0001.

To investigate regional differences in the association of osteocytes with different vascular sources, distance maps quantified lacunar organisation in two ways: 1) in relation to both bone surfaces and intracortical canals (Ca. Incl distance maps, Fig 3A) and 2) solely in relation to bone surfaces (Ca. Excl distance maps, Fig 3B). This enabled comparison of the minimum lacunar distance with (Ca. Incl) and without (Ca. Excl) intracortical canals, thus identifying populations of osteocyte lacunae closely associated with intracortical canals and those which would be isolated should intracortical canals not be present.

Global analysis of Ca.Excl distance maps compared with Ca. Incl maps show that intracortical canals reduce global Mm.Lc.D from 48.8 µm (± 1.9 µm) to 36.23 µm (± 0.9, p = 0.002). This finding was specific to the posterior region with intracortical canal inclusion in distance maps reducing Mm.Lc.D from 60.9 µm (± 6.4 µm) in Ca. Excl distance maps to 34.2 µm (± 1.8 µm) in Ca.Incl distance maps (p < 0.0001). For the anterior (p = 0.346), lateral (p = 0.81) and medial (p = 0.19) regions canal inclusion no significant effect was detected (Fig. 3C). Further quantification of the number of isolated lacunae was completed through calculation of the proportion of lacunae found > 100 µm from a bone surface. In the posterior region in Ca. Excl maps 26.8% (± 2.9%) were > 100 µm from a potential vascular supply, while Ca. Incl greatly reduced this (3.4% ± 0.6%, p < 0.0001, Data not shown).

### Canal-associated lacunae are the largest in **the** posterior region

To determine whether the abundant intracortical canals in the posterior TFJ support the large osteocyte lacunae in this region, lacunar populations were separated into canal-associated and surface-associated lacunae as illustrated in Fig. 1C. The posterior TFJ had more canal-associated lacunae (42.4% ± 4.5% of total lacunar number) than the anterior (20.1% ± 4.6%, p = 0.007), lateral (21.0% ± 1.2%, p = 0.009), and medial (20.9% ± 7.0%, p = 0.009) regions (Fig. 4B).

**Figure 4.**
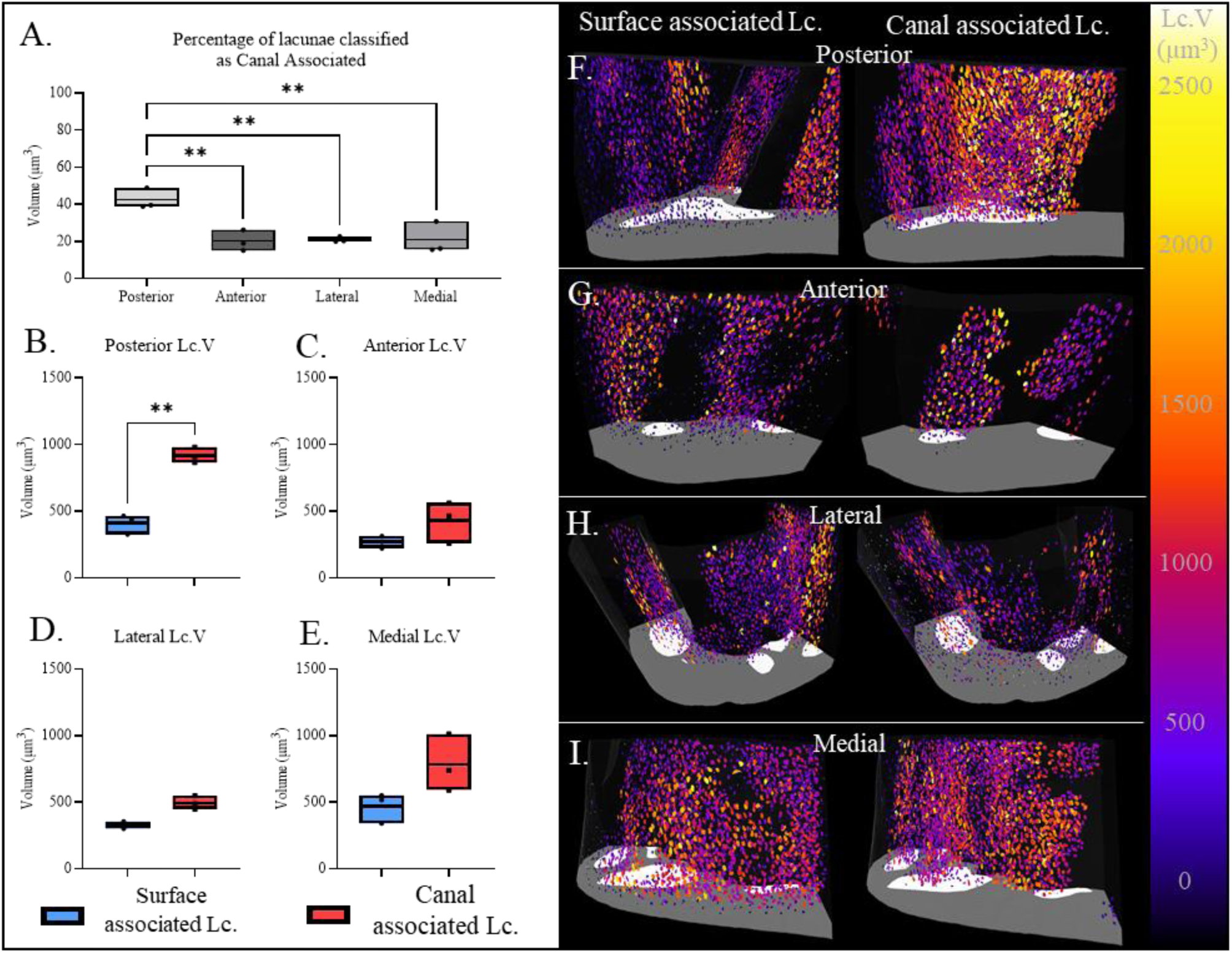
Lacunae associated with intracortical canals are larger than surface lacunae in the posterior region. (A) Quantifies the proportion of lacunae defined as canal-associated within each region, with more lacunae associated with intracortical canals in the posterior region versus other regions. The mean volume of canal and surface-associated lacunae was calculated in the posterior (B), anterior (C), lateral (D) and medial (E) regions. 3D visualisation of canal and surface-associated osteocyte lacunae volume distributions in the posterior (F), anterior (G), lateral (H) and medial regions (L). Larger lacunae are coloured in orange/yellow and smaller in blue/purple. Data is shown as mean ± SD and assessed by a one-way ANOVA with subsequent post hoc analysis. * Denotes p<0.05, ** p < 0.01, *** p <0.001.

In the posterior region canal-associated lacunae (919 µm^3^ ± 48 µm^3^) were larger than surface-associated lacunae (411 µm^3^ ± 60 µm^3^, p < 0.001, Fig. 3D). In the anterior (Fig. 4E), lateral (Fig. 4F) and medial (Fig. 4G) regions, no significant difference was found in the average lacunar volume between canal-associated and surface-associated lacunae. Further, posterior canal-associated lacunae (919 µm^3^ ± 48 µm^3^) were larger than those in the anterior (429 µm^3^ ± 126 µm^3^, p = 0.010) and lateral regions (494 µm^3^ ± 42 µm^3^, p = 0.022).

### Effect of VEGF deletion on vascular: lacunar spatial arrangements

To study the nature of lacunar-vascular associations, the OcnVEGFKO murine model was utilised. Disruption of VEGFKO in male mice leads to a highly vascularised TFJ (Goring et al. 2019), while here, male OcnVEGFKO cortices were similarly highly porous (Fig. 5A-B) and heavily vascularised (Fig. 5C) with no significant changes to mean osteocyte lacunar volume (Fig. 5D).

**Figure 5.**
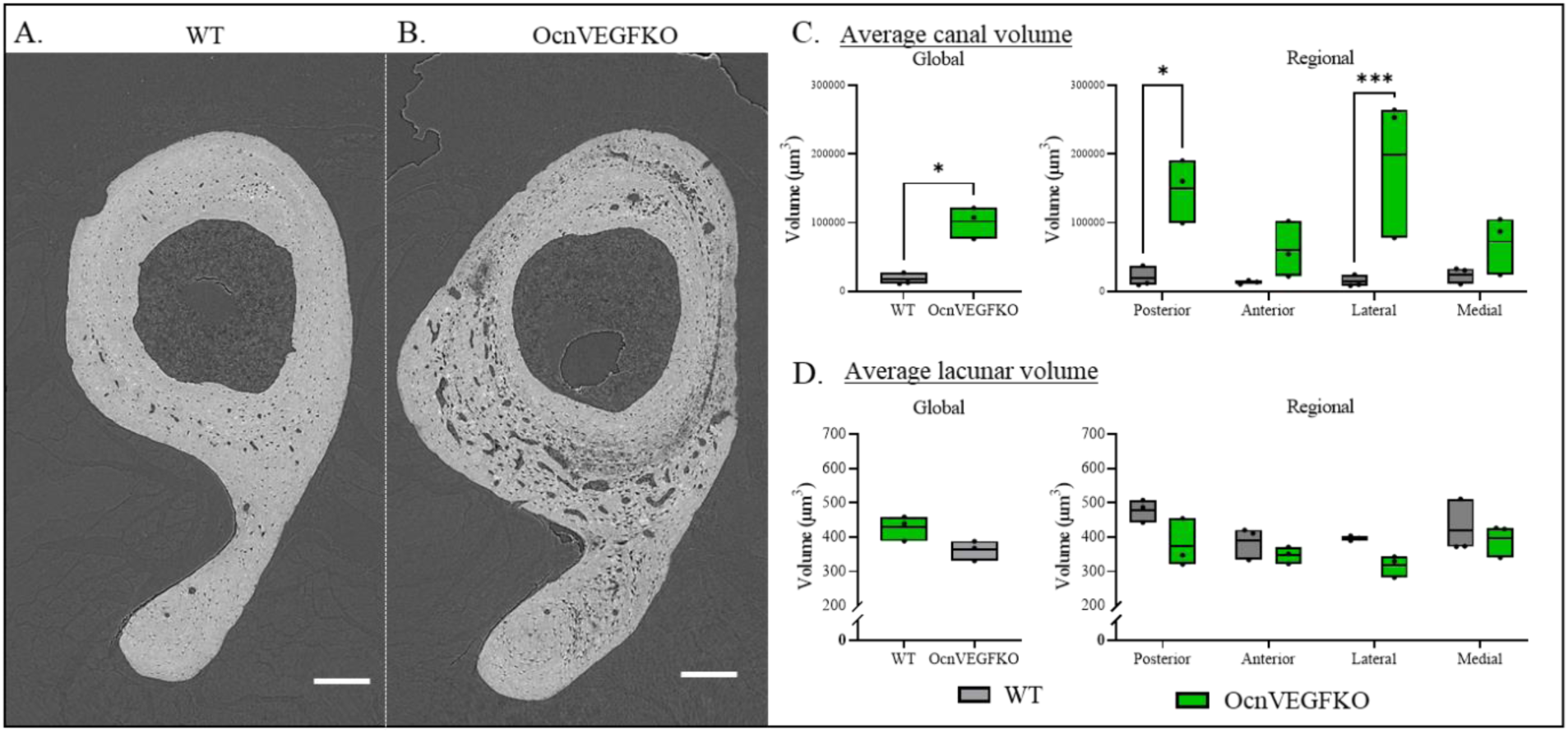
Conditional VEGFKO deletion disrupts intracortical canal volume in a region-specific manner. SR CT visualises male WT (A) and OcnVEGFKO (B) cortical bone at the TFJ, scale bar = 200μm. OcnVEGFKO mice possess larger intracortical canals (C) which upon regional assessment is localised to the posterior and lateral TFJ. Average lacunar volume is unaffected in OcnVEGFKO bone (D). N=3 WT and OcnVEGFKO, with littermates matched through paired comparisons. Data is presented as mean ± SD *p < 0.05, ***p < 0.001 using two-way ANOVA.

Osteocyte lacunae were separated into canal-associated and surface-associated in WT (Fig 6A-B) and OcnVEGFKO bone (Fig. 6C-D). There were no differences in the volume of surface-associated or canal-associated lacunae volume. Regionalisation of bone cortices revealed that in WT bone, posterior canal-associated lacunae (564 µm^3^ ± 42 µm^3^) were larger than surface-associated lacunae (439 µm^3^ ± 11 µm^3^, p = 0.034). Meanwhile, in OcnVEGFKO bone, we did not find any significant difference in average volume between canal-associated lacunae (346 µm^3^ ± 49 µm^3^) and surface-associated lacunae (342 µm^3^ ± 22 µm^3^, p = 0.120, Fig. 6E), suggesting that loss of VEGF signalling by osteocalcin expressing cells prevents the formation of distinct, large lacunar populations.

**Figure 6.**
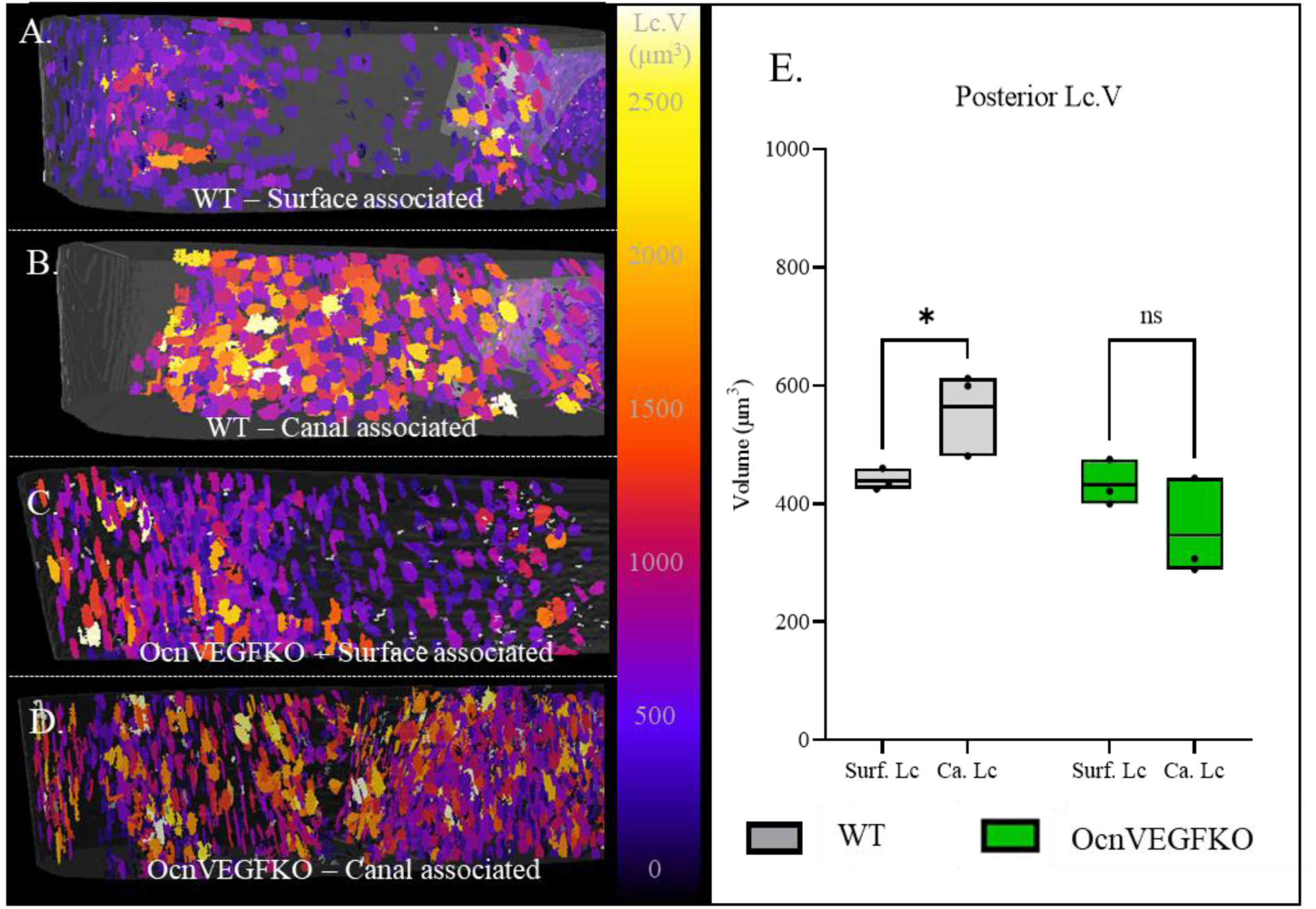
Conditional VEGFKO deletion leads to a homogenous population of osteocyte lacunae in the posterior region. 3D visualisation of surface-associated (A) and canal-associated (B) osteocyte lacunae within the posterior TFJ of 16-week-old male mice WT mice (n=3) and OcnVEGFKO mice (C.D respectively) with large lacunae shown in yellow and white and small lacunae in purple. (E) Quantification of average Lc.V between WT and OcnVEGFKO mice. Data is shown as mean ± SD and assessed by a two-way ANOVA with subsequent post-hoc analysis. * Denotes p<0.05, ** p < 0.01, *** p <0.001.

**Figure 7.**
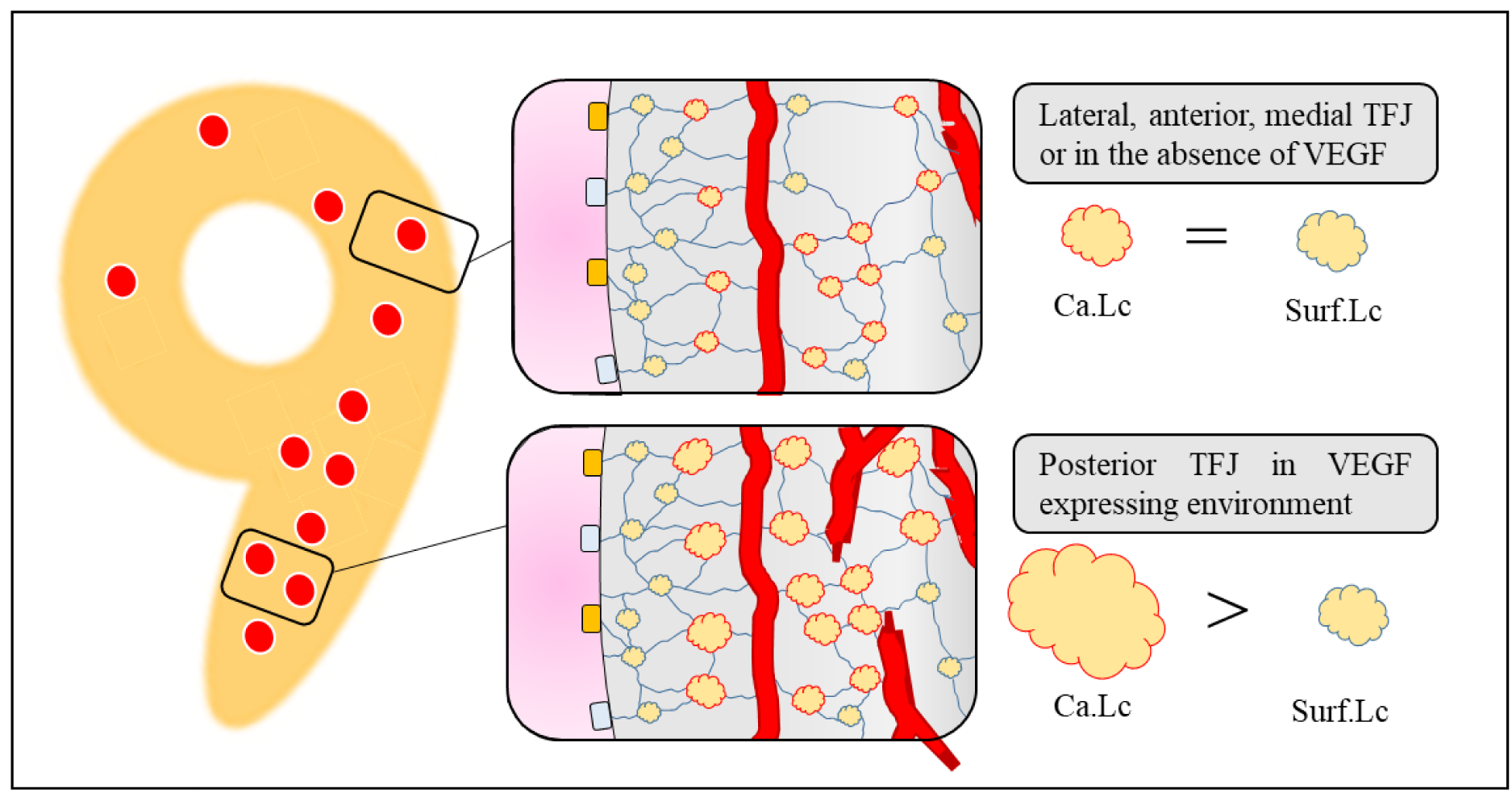
Canal proximity within the posterior TFJ supports large osteocyte lacunae. WT mice possess larger canal-associated lacunae than surface-associated lacunae, specific to the posterior TFJ with canal proximity not relating to changes in lacunar volume in anterior, lateral and medial regions. In contrast, in OcnVEGFKO bone canal-associated and surface-associated osteocyte lacunae at the posterior TFJ are homogenous in volume. Disruption of canal-associated lacunar volume by VEGFKO could be indicative of the role of VEGF in the direct regulation of the osteocyte lacunae network and its interactions with the bone vasculature.

## Discussion

This study provides evidence that the cortical microstructure is heterogeneous and that a population of large osteocyte lacunae surround intracortical canals in a region-specific, VEGF-dependent manner. Upon development of an automated regionalisation tool, we identify the posterior region of the TFJ as a site possessing abundant intracortical canals and large osteocyte lacunae. Mapping of the spatial organisation of osteocyte lacunae surrounding intracortical canals revealed that large lacunae surround the abundant intracortical canals within the posterior region. Notably, this distinct population of canal-associated, large lacunae did not form in a murine model of conditional VEGF-knockout at the posterior TFJ.

Previously, regional assessment of cortical bone has been used to describe region-specific alterations to the bone cortex utilising both 2D (Cole et al. 2022; van Tol et al. 2020) and 3D imaging modalities (Núñez et al. 2018; Uniyal et al. 2021; Schneider et al. 2007; Chiba et al. 2013). However, the use of manual segmentation for regional selection lacks reproducibility and thus consistency between studies. This work describes an automated technique in open-source software that enables accurate and reproducible regionalisation of the murine bone cortex, applicable to large 3D datasets. Our proposed method has been applied to separate the C57BL/6 TFJ into posterior, anterior, lateral, and medial regions. Prior regional assessment of bone has confirmed that specific cortical regions are prone to deterioration (Núñez et al. 2018; van Tol et al. 2020), with the posterior TFJ susceptible to the effects of ageing (Núñez et al. 2018). This ability to regionalise the cortex permits the combination of techniques such as SR CT and finite element analysis (FEA) (Núñez et al. 2018; Zahm et al. 2010) to assess the effects of loading (Donaldson et al. 2014; Larrue et al. 2011) and the effects of microstructural deterioration (van Tol et al. 2020) on cortical bone’s susceptibility to fracture. However, as this method is developed for the murine TFJ, further validation is required to determine its applicability in investigating regional-specific functions of the bone microstructure at other anatomical sites.

Regionalisation of the TFJ revealed that the posterior TFJ possessed a greater number density of intracortical canals versus the other regions of the TFJ, concordant with the literature (Núñez et al. 2018; Maeda et al. 2020). The finding that intracortical canal distribution is heterogeneous is not specific to the murine TFJ however: at the murine femoral midshaft, the lateral region possesses more canals than the posterior region (Schneider et al. 2007) while in humans the medial proximal femoral shaft houses more intracortical canals than the anterior and lateral regions (Perilli et al. 2015). This may suggest that the abundance of intracortical vasculature may vary between specific anatomical sites; however, what links these sites, and the underpinning vascular heterogeneity remains unspecified. One hypothesis is that mechanical strain may govern the formation of intracortical canals: the inhomogeneous distribution of mechanical strain at the murine TFJ, and the relative absence of mechanical strain in the posterior region (Núñez et al. 2018), could suggest that mechanical strain negatively regulates the formation of the bone vasculature. Alternatively, whether these abundant intracortical canals within the posterior TFJ provide a specific function is unknown. One suggestion is that the posterior TFJ is a region of thicker bone cortex (Núñez et al. 2018), suggesting that such intracortical canals deep within the posterior cortex may prevent formation of a hypoxic microenvironment to sustain populations of lacunae deep within the bone cortex. Here, we report that large osteocyte lacunae are located around this distinct population of abundant posterior intracortical canals, suggesting that this subset of canals may provide a specific, osteogenic function tied to large osteocyte lacunae.

Variability in osteocyte lacunae volume has been observed in healthy murine (Núñez et al. 2018) and human bone (Carter et al. 2013), as well as across taxa including ray-finned fishes (Davesne et al. 2020), suggesting that variability in osteocyte lacunae volume is a conserved feature for the maintenance of cortical bone. Further, in diseased human bone, osteopenic bone is characterised by large osteocytes while osteoarthritic bone is populated by small osteocytes (van Hove et al. 2009). The function of osteocyte lacunae heterogeneity – in particular, large lacunae associating with the bone vasculature within specific bone regions - remains unclear, with hypotheses primarily focused on the potential role of the LCN in mechanotransduction (Schurman, Verbruggen, and Alliston 2021; Qing et al. 2012; Lane et al. 2006).

A dampened mechanoresponse in aged mice has been reported to be facilitated by reduced LCN connectivity which interestingly, is rescued upon expansion of the lacunar space surrounding the osteocyte cell body, suggestive that enlargement of the pericellular space may be compensatory in the response to a reduced LCN connectivity (Schurman, Verbruggen, and Alliston 2021). While the potential for a heterogeneous distribution of large osteocyte lacunae playing differential roles in strain distribution requires further interrogation, the reporting of heterogeneity in osteocyte canalicular length and load-induced fluid flow at the murine TFJ (van Tol et al. 2020) does suggest that these adaptions to LCN degeneration may be region-specific. This, in tandem with the finding that the posterior TFJ is susceptible to microstructural degeneration (Núñez et al. 2018), highlights this region as particularly prone to degeneration and as such, the presence of large osteocyte lacunae may be indicative of a larger LCN response to microstructural degeneration.

Alternatively, the role of osteocytes in bone (re)modelling (Bonewald 2011; Kennedy et al. 2014) could suggest that the heterogeneous distribution of large osteocyte lacunae facilitates regional bone formation – such as during perilacunar remodelling (PLR). Regulation of lacunar and canalicular volume through local matrix remodelling is observed in response to lactation (Qing et al. 2012) and glucocorticoid treatment (Lane et al. 2006). Therefore, the population of large lacunae at the posterior TFJ region may be lacunae currently resorbing the surrounding matrix, increasing their lacunar volume. Should large lacunae differentially affect the composition of the local matrix versus small lacunae in specific regions of bone, their potential for the regulation of bone mineral may provide avenues for the research of region-specific bone anabolic agents.

However, the correlative appearance of abundant intracortical canals and large lacunae may suggest a paired functionality. Assessment of the regional spatial organisation of osteocyte lacunae in relation to intracortical canals in this study found that posterior intracortical canals prevent the isolation of a subset of large lacunae. This may suggest that preferential vascular support of large lacunae by intracortical canals is a region-dependent phenomenon, which may facilitate region-specific functionality, highlighting the candidacy of the posterior region of the TFJ as a site of unique angio: osteogenic coupling. Heterogeneity in vascular functionality and the facilitation of osteogenesis is well described (Ramasamy et al. 2014; Kusumbe, Ramasamy, and Adams 2014; Ramasamy et al. 2016), with a specific vascular subtype - type H blood vessels - shown to be a key regulator of osteogenesis. Type H vessels have been shown to upregulate osteoprogenitor survival and proliferation factors through Notch (Ramasamy et al. 2014) and HIF1α (Kusumbe, Ramasamy, and Adams 2014) signalling. However, how Notch signalling regulates the volume of the LCN is unknown, with Notch signalling in osteocytes inducing an osteopetrotic (high bone mass) phenotype (Canalis et al., 2015) – a condition associated with small lacunae (van Hove et al. 2009). Furthering our understanding of how specific vascular subsets regulate osteocyte volume and the regulation of PLR may elucidate how and why these large lacunae exist.

Thereby, we hypothesise that the removal of factors coupling angiogenesis and osteogenesis would prevent the formation of large osteocyte lacunae surrounding posterior intracortical canals. Previously, induction of hype-H endothelial cell formation has been shown to elevate bone marrow and serum levels of VEGF (Xie et al. 2014). Furthermore, osteocyte-like cell lines express VEGF in response to shear stress placed on canaliculi (Harper, Gerstenfeld, and Klagsbrun 2001) and hypoxia (Mulcrone et al. 2020) which may suggest that in response to mechanical and hypoxic stimuli, osteocytes have the potential to regulate angiogenesis.

As such, the organisation of osteocyte lacunae was assessed in a murine model with conditional VEGF deletion in osteocalcin-expressing cells (Goring et al. 2019). Here, we show that at the posterior TFJ of male OcnVEGFKO mice, lacunar volume was unaffected by proximity to the intracortical vasculature, in any region. Thereby, loss of VEGF prevents the formation of this region-specific, population of large lacunae.

While the effects of VEGF deletion have been described(Goring et al. 2019) and its regulation of the bone microstructure, with male OcnVEGFKO bone characterised by mineralisation deficits and increased SOST expression (Goring et al. 2019), suggesting that loss of VEGF may affect osteocyte function, which in tandem with an apparent inability to form this sub-population of large osteocyte lacunae in the posterior region of the TFJ, may suggest that these large osteocytes play a role in the regulation of bone mineralisation. As such, moving forward, assessment of the mineral surrounding canal-associated osteocytes in health, disease and treatment states could provide insight into the potentially unique functionality of these canal-associated osteocytes. Currently, techniques that enable the 3D assessment of osteocytes within their lacunae and subsequent relation to the mineral matrix (i.e., without decalcification) are limited, with the majority of applications lacking sufficient soft tissue contrast and resolution to visualise the osteocytes within lacunae.

The identification of a population of large osteocyte lacunae closely associated with densely vascularised cortical regions such as the posterior TFJ, suggests that within specific regions of the bone cortex, there may be regions utilising unique osteo: angiogenic-coupling mechanisms that may underpin the regulation of osteocyte lacunae volume. Furthermore, we show that in the absence of the key angiogenic factor, VEGF, these large osteocyte lacunae do not form at the posterior TFJ, suggestive of uncoupling of this region-specific osteo: angiogenic signalling.

## Acknowledgements

This study was jointly funded by the Gerald Kerkut Charitable Trust and the University Of Southampton. We would like to acknowledge the team at the IL-13 Beamline at Diamond Light Source and the µ-VIS X-ray Imaging Centre at the University Of Southampton for providing access to image analysis PCs. We would also like to thank the team at Object Research Systems (ORS) for providing a non-commercial license for the use of Dragonfly.

## Authors’ roles

Study design: JT, AS, PG, CEC. Data collection: JT, AS. Data analysis JT. Data interpretation: JT, CEC. Drafting manuscript: JT, PS, and CEC. JT takes responsibility for the integrity of the data analysis.

